# A NOVEL APPROACH TO RECOVERING OFF-TRACK ALIGNER TREATMENT THROUGH HYPER-ELASTIC POLYMER INNOVATION

**DOI:** 10.1101/2024.03.25.586642

**Authors:** Loc X. Phan, Carla Mora, Agnieszka Klucinska

## Abstract

Clear plastic aligners are becoming an increasingly popular choice for patients seeking orthodontic treatment. However, Industry Standard Aligners lack material properties that sustain tooth movement throughout the typical wear time for aligners.

In a study compiling force data from existing Industry Standard Aligners and the OrthoFX Rescue Aligner, data points were plotted to compare the force profiles of each product. The Rescue Aligner aims to eliminate the mid-course correction process and provide consistent forces to improve treatment times and patient satisfaction.

## Introduction

Clear plastic aligners have become a common and preferred orthodontic treatment method over time, due to their convenience and easy concealment. However, with the limitations of the semi-flexible material typically used in clear plastic aligners, deformations occur that can lead to ineffective tooth movement and time during treatment in which no progress is made. This material makes it increasingly difficult to deliver a constant force necessary for orthodontic tooth movements (OTMs), as it suffers from stress decay due to a process called viscoelastic hysteresis.

OrthoFX has introduced an innovative treatment option called Rescue Aligners. Advanced beyond the conventional semi-flexible materials, the Rescue Aligner has hyper-elastic properties that provide constant forces during the course of its 7-day wear time. This new property of “hyper-elasticity” avoids the steep force profile drop offs plaguing Industry Standard Aligners.

Treatment deviation requires mid-course correction, which typically takes six to eight weeks and multiple office visits before correction begins. The OrthoFX Rescue Aligner eliminates the mid-course correction process and associated appointments altogether by shipping a Rescue Aligner directly to the patient to begin correction within one week of detection. Combining a dramatically reduced correction process with the hyper-elastic material that yields a consistent forcfie profile throughout OTM, Rescue Aligners provide an overall faster and more efficient solution to treatment deviations.

## Material and Methods

To adequately compare an OrthoFX Rescue Aligner to an Industry Standard Aligner, it is necessary to determine the effectiveness of the aligners by measuring the force produced by each aligner throughout the desired movement period. This was achieved through the compilation of data from multiple angular force investigations, specifically ±3° of Upper Right Canine Torque and Rotation as well as a linear force investigation compiling data from ±0.3mm translation of an Upper Right Lateral Incisor Expansion/Contraction. These OTM’s were chosen based on the difficulty of standard industry aligners to provide consistent force throughout treatment *(Fig. 1)*. As seen in *Figures 2 through 8*, OrthoFX collected 7,680 data points through a variety of OTM treatments to best compare the force outputs produced by an Industry Standard Aligner and an OrthoFX Rescue Aligner.

**Figure 1:**
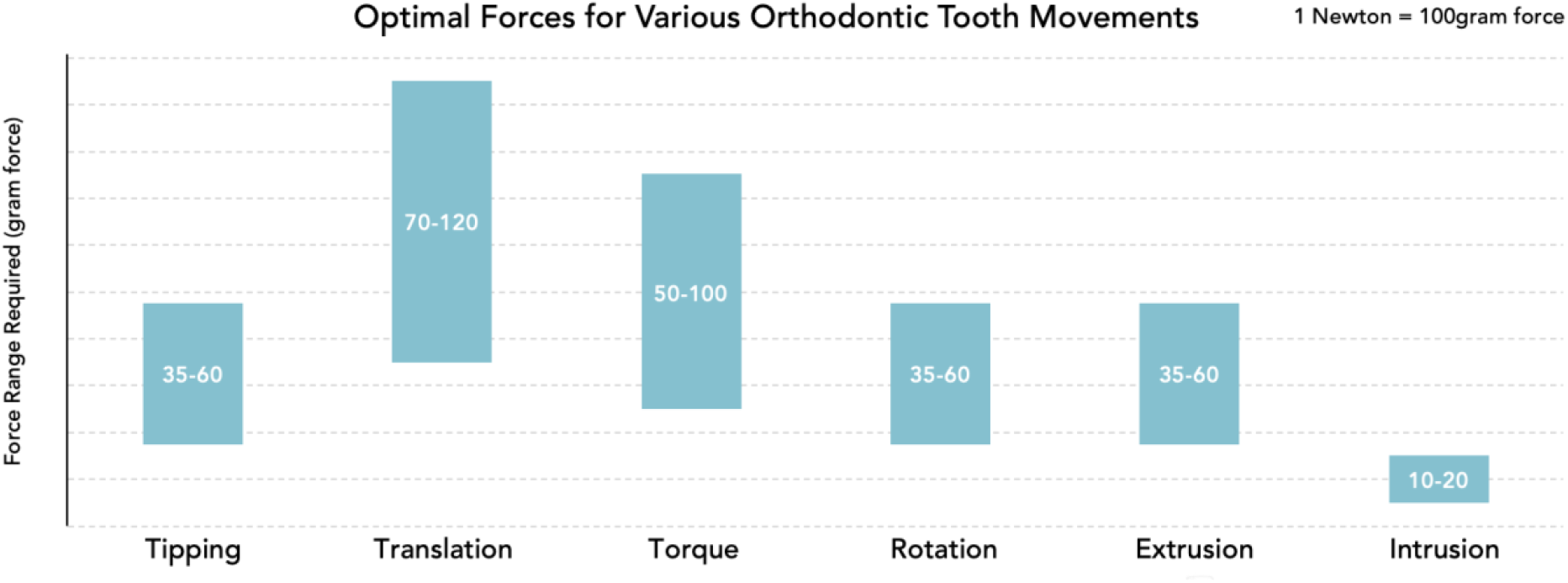
Range of force typically needed to achieve various OTMs.

**Figure 2:**
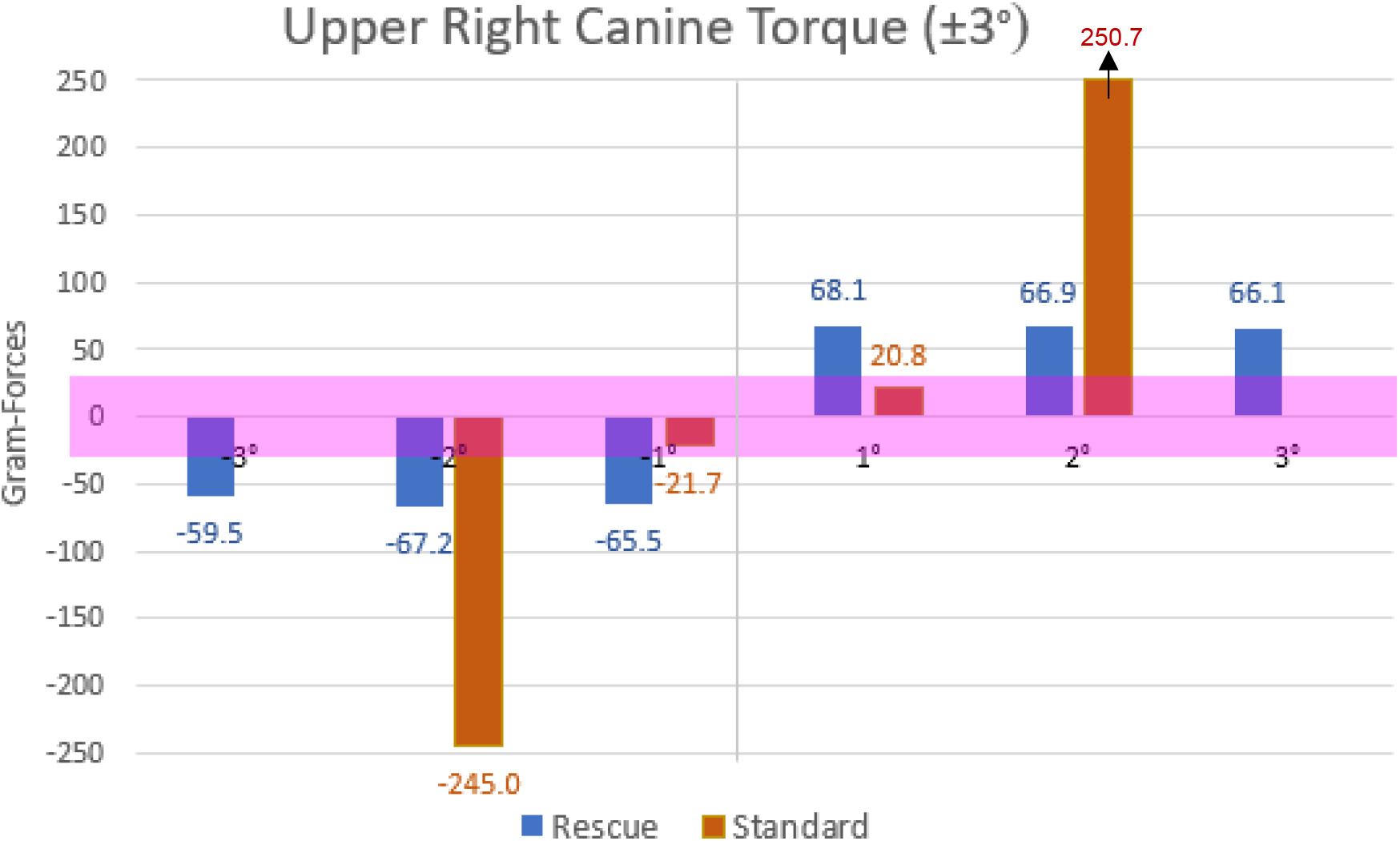
Force produced by Industry Standard Aligner vs. OrthoFX Rescue Aligner throughout a ±3° Torque OTM.

## Results

### Angular Force Data

A comparison of ±3° of Upper Right Canine Torque using a Standard Industry Aligner versus an OrthoFX Rescue Aligner is plotted in *Figure 2*. A purple bar representing the ideal biological force range is plotted at ±25 grams of force. A Standard Industry Aligner provides -21.7 and 20.8 grams of force within the first degree of rotation followed by a steep increase to -245.0 and 250.7 grams of force for the second degree and failed to provide any force for the final third degree of torque. The OrthoFX Rescue Aligner provides a consistent force range varying between -59.5 to -67.2 grams and 68.1 to 66.1 grams for all three degrees of torque.

Similar to the torque comparison, a ±3° of Upper Right Canine Rotation using a Standard Industry Aligner versus an OrthoFX Rescue Aligner is plotted in *Figure 3*. A purple bar representing the ideal biological force range is plotted at ±25 grams of force. The Standard Industry Aligner provides - 14.6 and 15.4 grams of force within the first degree of rotation followed by a steep increase to -246.3 and 201.0 grams of force for the second degree and failed to provide any force for the final third degree of rotation. The OrthoFX Rescue Aligner provides a consistent force range varying between -34.9 to -35.6 grams and 34.8 to 35.1 grams for all three degrees of rotation.

**Figure 3:**
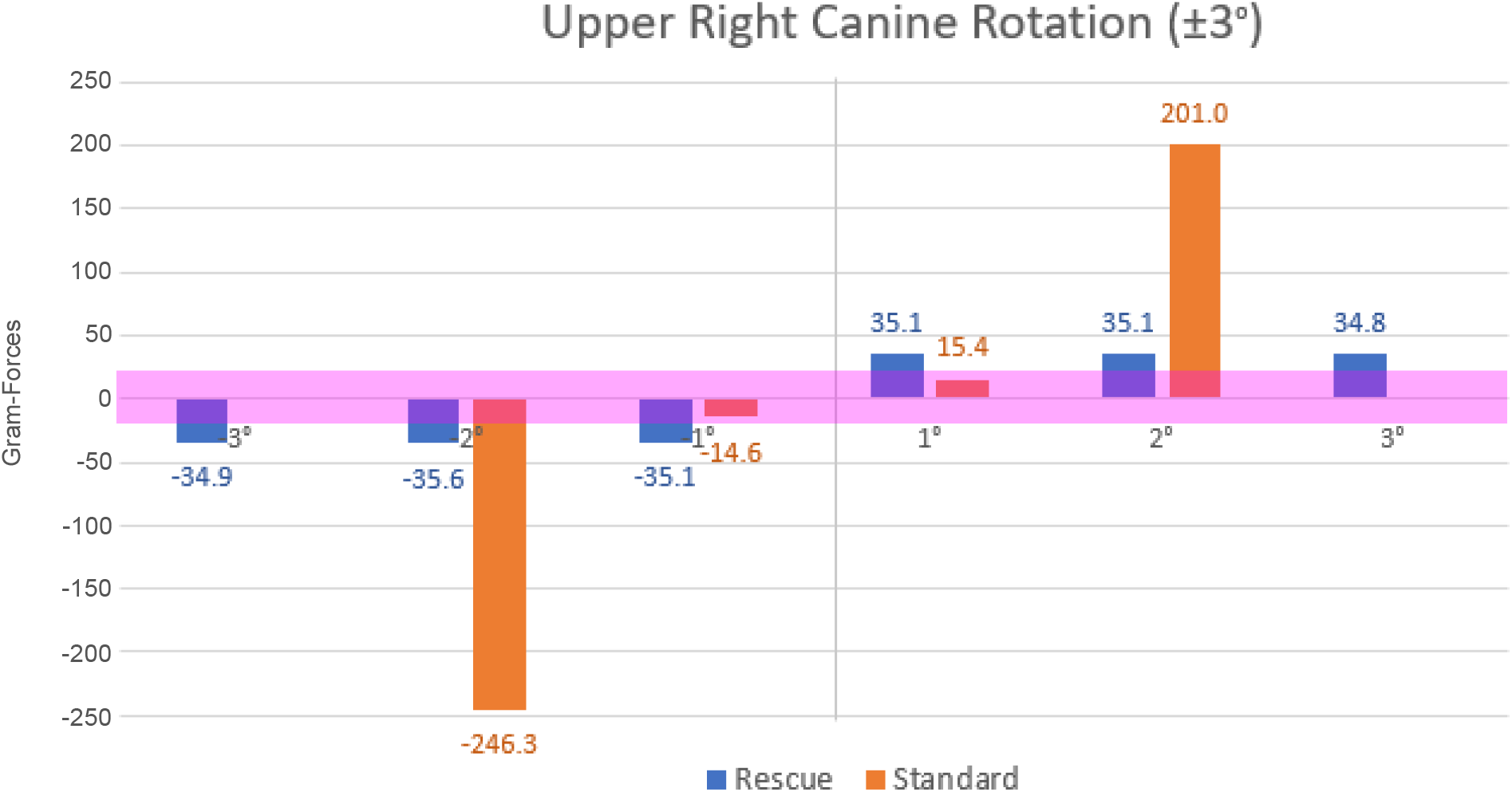
Force produced by Industry Standard Aligner vs. OrthoFX Rescue Aligner throughout a ±3° Rotation OTM.

### Linear Force Data

A comparison of ±0.3mm of Upper Right Lateral Incisor Expansion/Contraction using a Standard Industry Aligner versus an OrthoFX Rescue Aligner is plotted in *Figure 4*. A purple bar representing the ideal biological force range is plotted at ±70 grams of force required for linear movement. The Standard Industry Aligner provides -55 and 54 grams of force within the first 0.1mm of translation, followed by a steep increase to -282 and 355 grams of force at 0.2mm of translation and failed to provide any force at 0.3mm of translation. The OrthoFX Rescue Aligner provides a consistent force range varying between - 96 to -101 grams and 96 to 99 grams for all 0.3mm of translation.

**Figure 4:**
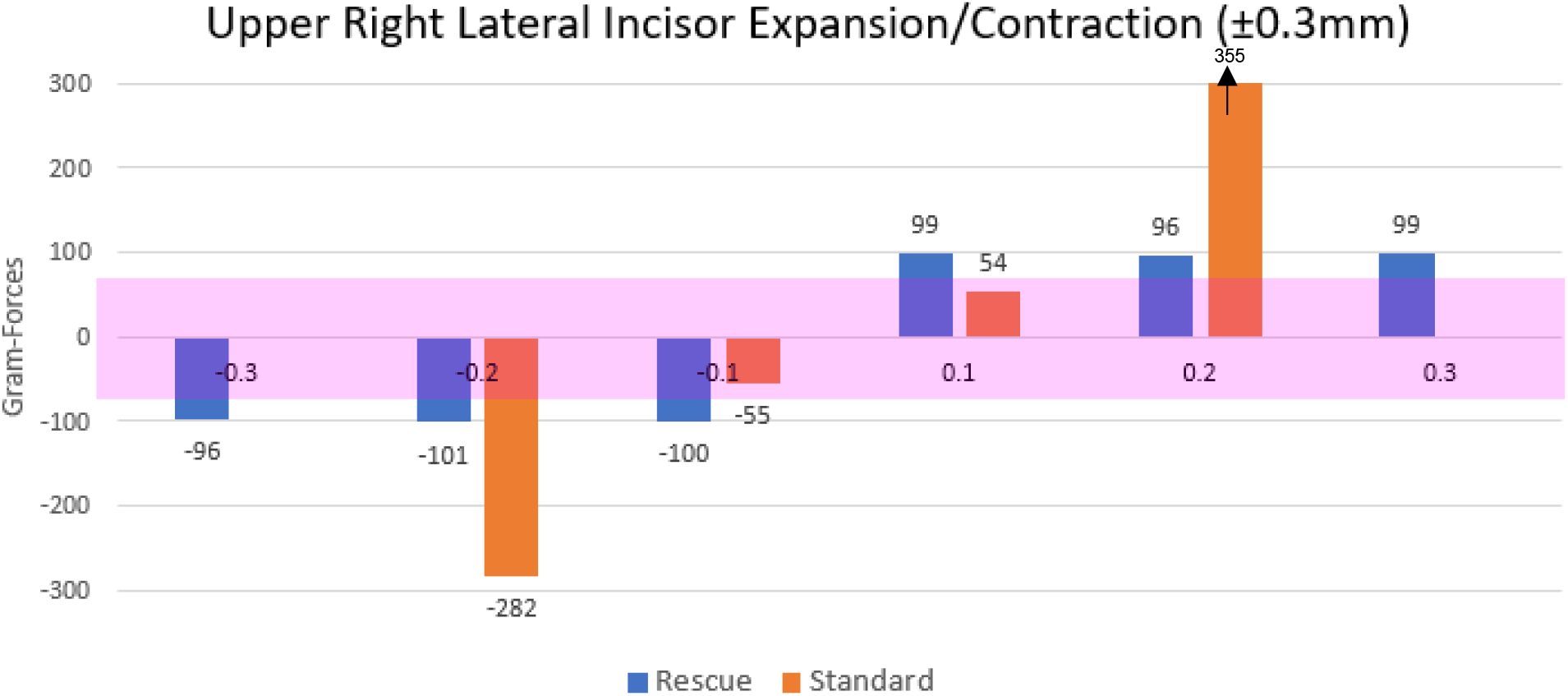
Force produced by Industry Standard Aligner vs. OrthoFX Rescue Aligner throughout a ±0.3mm Translation OTM.

### Force Profile Data

When comparing force profiles of the OrthoFX Rescue Aligner (*Fig. 6*) versus Standard Industry Aligners (*Fig. 5*), it is important to note the initial force exerted by the aligner. While a Standard Industry Aligner exerts 800 grams of force initially, the OrthoFX Aligner only exerts roughly a sixth of this force at 140 grams of force. However, note in *Figures 5 and 7*, the abrupt decline of the Standard Industry Aligner’s high initial force within the first five hours of use. This decline then continues into the second day of use, falling below the desired optimal force level and never recovering. Conversely, in *Figures 6 and 8*, the OrthoFX Rescue Aligner does not suffer a steep decline, rather it drops to 100 grams at the end of the first day and finishes the second and fourth days at 80 grams of force or higher, which is still within the desired optimal force level.

**Figure 5:**
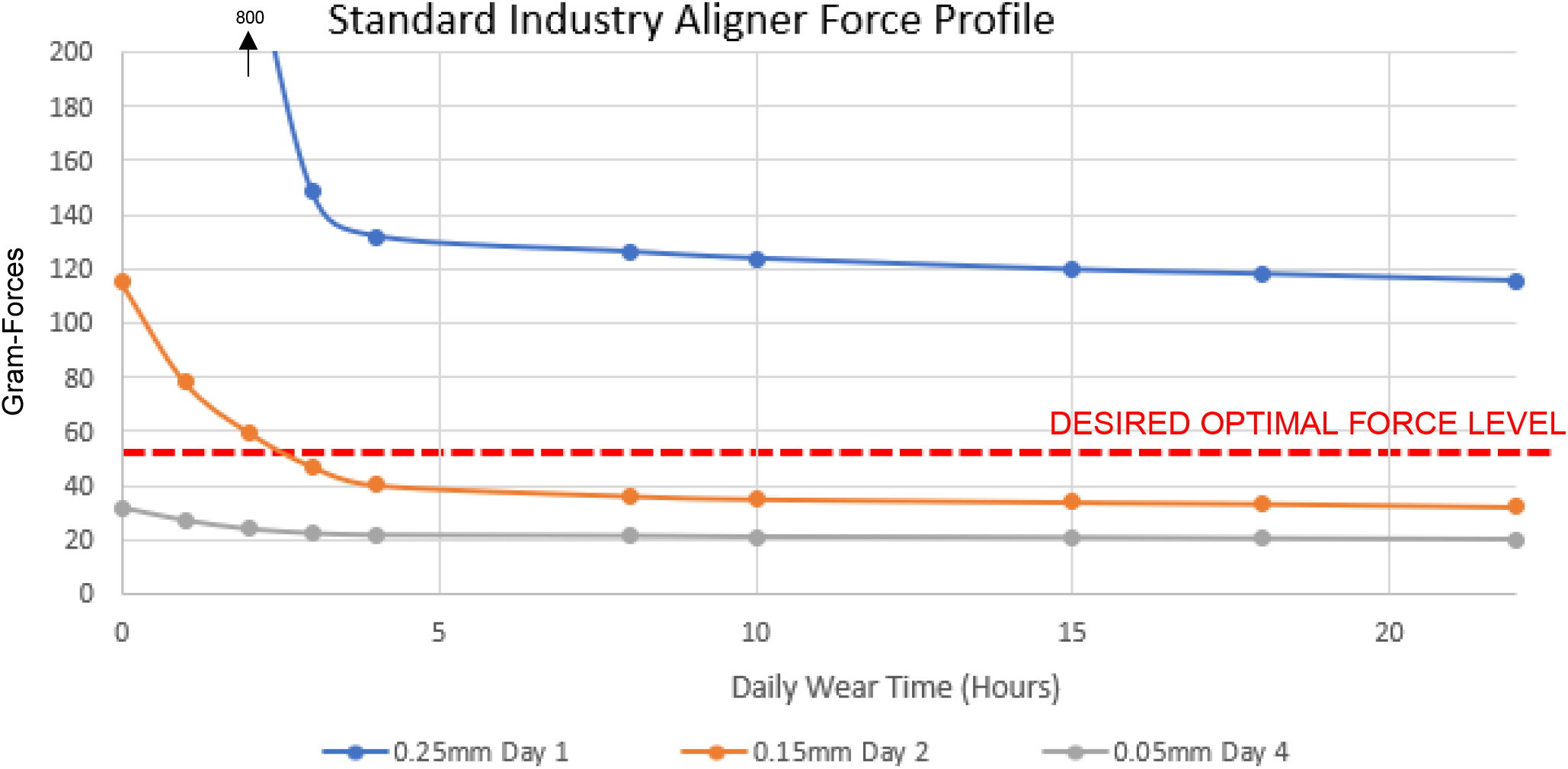
Force profile produced by Industry Standard Aligner throughout a 4-day period.

**Figure 6:**
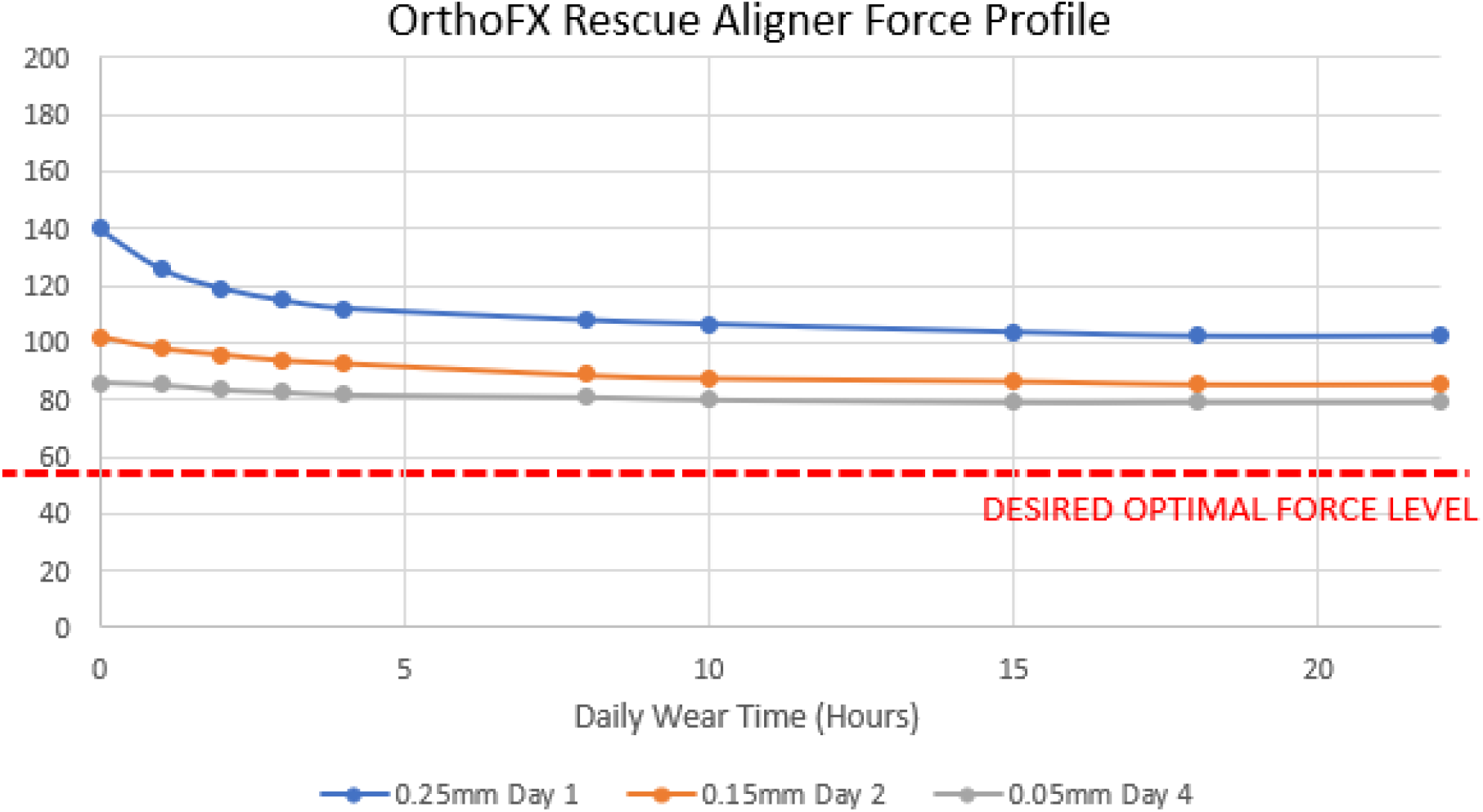
Force profile produced by OrthoFX Rescue Aligner throughout a 4-day period.

**Figure 7:**
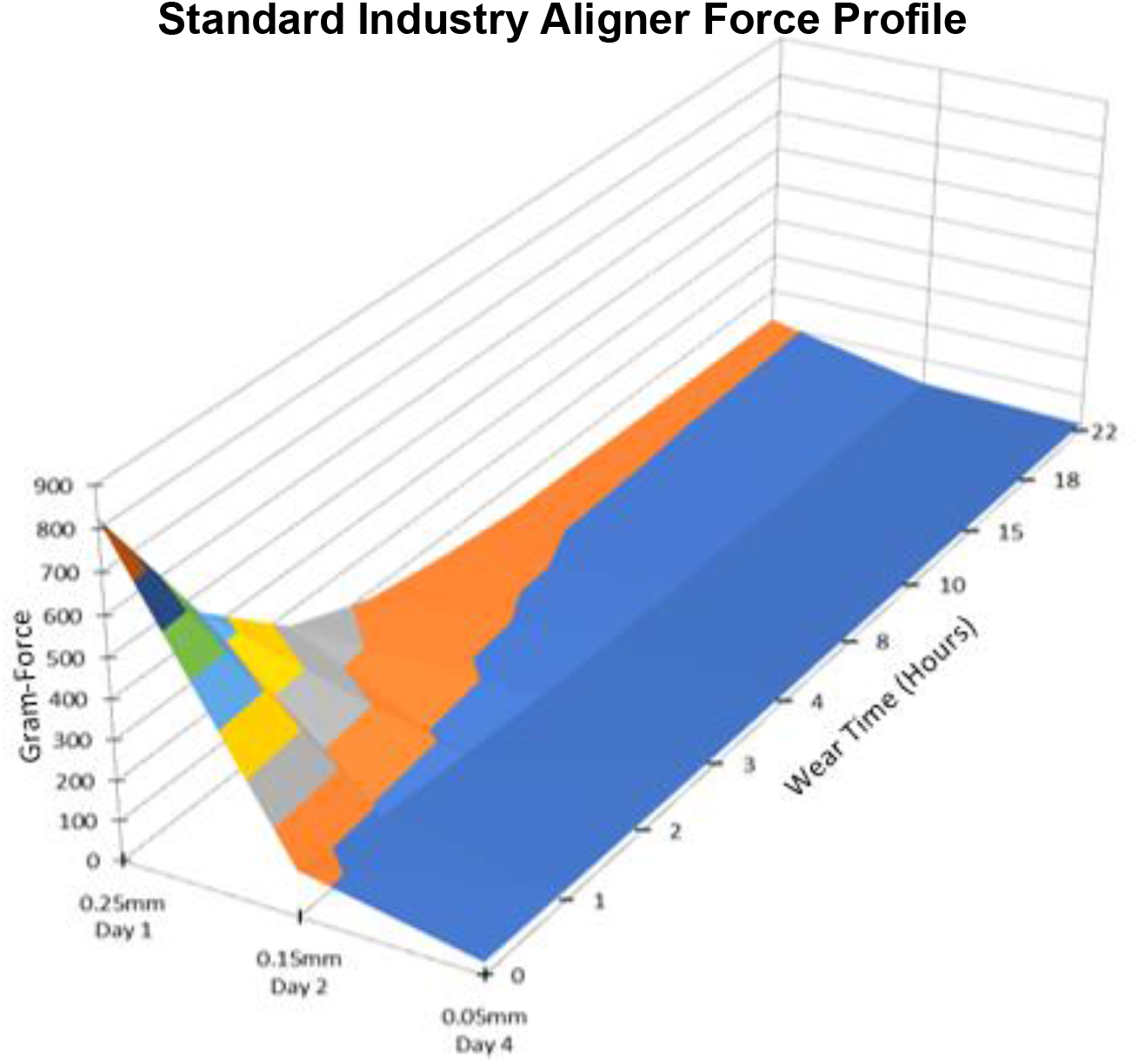
Force profile produced by Industry Standard Aligner throughout a 4-day period.

**Figure 8:**
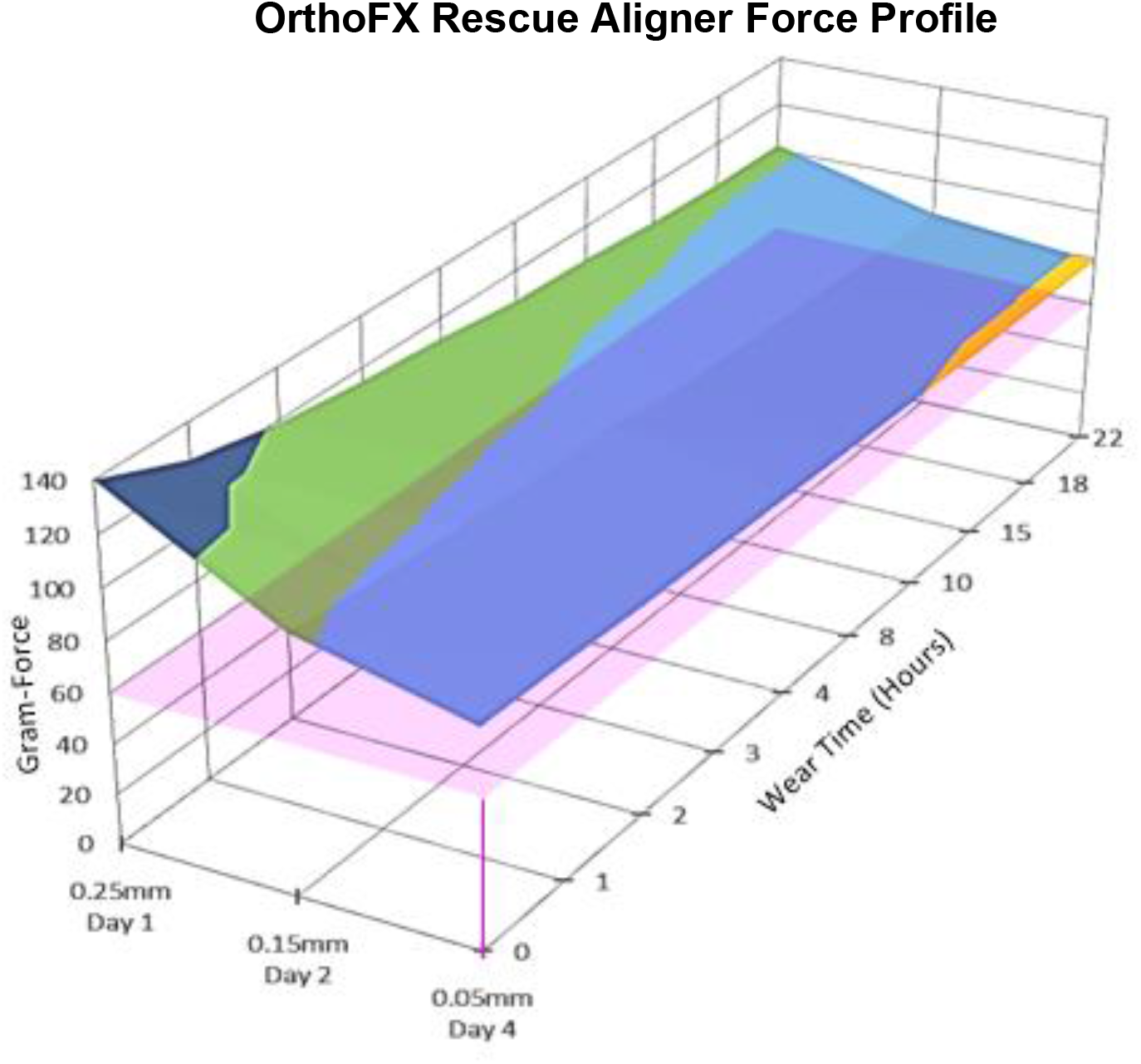
Force profile produced by OrthoFX Rescue Aligner throughout a 4-day period.

## Discussion

The OrthoFX Rescue Aligner provides consistent forces throughout a variety of OTM’s meant to test its capabilities against the Industry Standard Aligners. Typically starting with a high initial force that quickly tapers off once tooth movement begins, Standard Aligners provide an uncomfortable initial force for the patient and an ineffective use of time once the aligner is unable to support the desired OTM. This leads to the inconsistent forces displayed in *Figures 2-5, and 7* that progress from many times the desired force to suboptimal then quickly dissipating and providing no force whatsoever. This is due to the Standard Aligner suffering from viscoelastic hysteresis, a process involving energy dissipation following an applied load.

On the other hand, the OrthoFX Rescue Aligner begins and ends treatment in the desired range to encourage tooth movement, which evens force distribution over time, as seen in *Figures 2, 3, 4, 6, and 8*. This allows for a more comfortable experience for the patient and quicker OTM times. Due to the Rescue Aligner’s hyper-elastic material, the force required to continue an OTM is still available following increased strain. The Rescue Aligner comes as a clear-cut improvement to the current Industry Standard Aligner technology, increasing efficacy through consistent force outputs and saving patients significant time during treatment.

In addition to the hyper-elastic material advantages, Rescue Aligners also reduce treatment delays through a simpler and more straightforward mid-course correction process. Notice in *Figure 9*, the timesaving benefits of the OrthoFX Rescue Aligner’s approach. Office visits are no longer needed prior to implementing corrective action. With these improvements, the Rescue Aligner correction process cuts the standard wait time from six to eight weeks down to one week.

**Figure 9:**
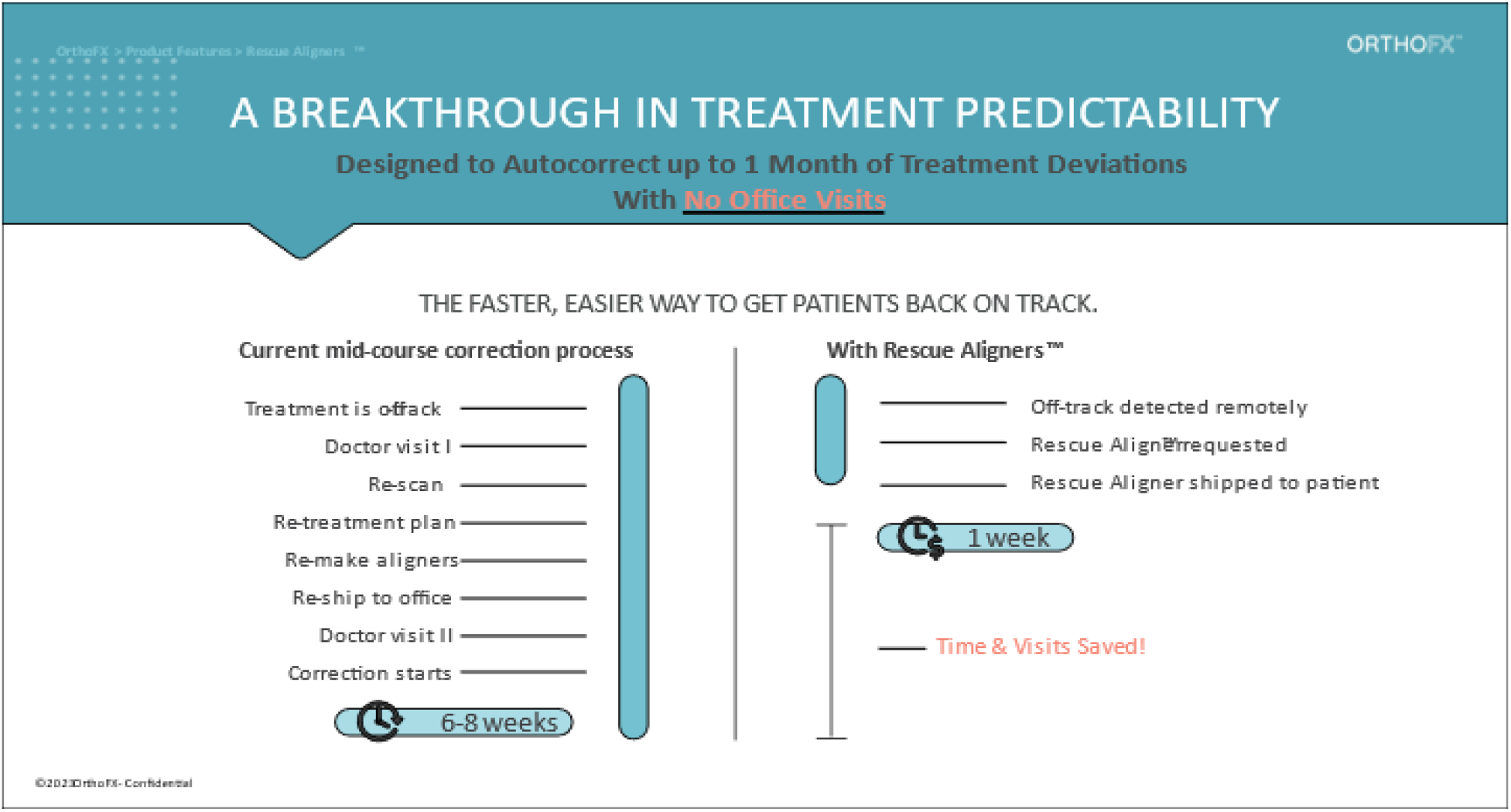
Industry Standard Aligner vs. OrthoFX Rescue Aligner treatment mid-course correction process.

## Conclusion

Through the compilation of test data comparing a variety of common OTMs such as torque, rotation, and translation, OrthoFX was able to compare its Rescue Aligner to Industry Standard Aligners. The results provided a comprehensive look at the force profiles produced by both aligners. Industry Standard Aligners start with a high force initially, then drop off before delivering planned tooth movement. The OrthoFX Rescue Aligner provides a consistent force profile that aims to achieve the biologically friendly level of force necessary to perform OTMs consistently in a minimal amount of time. The Rescue Aligner contributes to quality of care by improving upon both the convenience and efficiency of treatment.

## Notes

### Competing Interest Statement

All authors are employed at OrthoFX.

